# GEMC1 and CCNO are required for efferent duct development and male fertility

**DOI:** 10.1101/258418

**Authors:** Berta Terré, Michael Lewis, Gabriel Gil-Gómez, Travis H. Stracker

**Author notes:** To whom correspondence should be addressed. ORCID(0000-0002-8650-2081) Institute for Research in Biomedicine (IRB) Barcelona, Department of Oncology, C/ Baldiri Reixac 10, 08028 Barcelona, Spain.

## Abstract

GEMC1 is a Geminin family protein that triggers the E2F4/5-mediated transcriptional activation of target genes, including many required for multiciliogenesis, such as *Mcidas, FoxJ1* and *Ccno*. Male mice lacking either *Gemc1* or *Ccno* were found to be infertile, but the origin of this defect has remained unclear. Here we show that mice lacking either *Gemc1* or *Ccno* showed a nearly empty seminiferous tubule phenotype. While both genes were expressed in the testes, much higher levels were observed in the efferent ducts that mobilize sperm to the epididymis. We found that both GEMC1 and CCNO were required for the generation of multiciliated cells in the efferent ducts but that they differentially affect activation of *FoxJ1* and *Tp73*. These data indicate that defective efferent duct development, rather than defects in testes development, are likely the primary origin of male infertility observed in the absence of GEMC1 and CCNO and this could extend to Reduced Generation of Multiple Motile Cilia patients with *MCIDAS* and *CCNO* mutations.

**Summary statement:** We demonstrate that male mice lacking GEMC1 or CCNO are infertile due to defects in the formation of the efferent ducts.

## Introduction

Spermatogenesis is a highly regulated developmental process that generates haploid sperm capable of fertilization. It begins with the amplification of diploid spermatogonia through mitosis that is followed by meiosis to ensure the haploidization of recombined genomes. Finally haploid germ cells undergo terminal differentiation into mature spermatozoa, a process known as spermiogenesis. This is characterized by the remodeling of chromatin, reorganization of mitochondria and the development of the flagellum and acrosome. Eventually global transcription ceases, as the DNA condenses in the nucleus, and post-transcriptional events become crucial to regulate translation(Steger, 2001).

To become fully capable of fertilization, developed spermatids must detach from the seminiferous epithelium, enter the tubule lumen and travel through the epididymis that promotes maturation and motility of spermatozoa. The efferent duct (ED) connects the testes to the epididymis and it is important for sperm concentration, as it reabsorbs part of the luminal seminiferous fluid. The epithelia of the EDs contain poorly characterized multiciliated cells (MCCs) that mobilize the luminal fluids through the concerted action of motile cilia(Joseph et al., 2011).

Differentiation of MCCs in the airway is initiated in part through the action of the *Mir449* family of miRNAs(Kyrousi et al., 2015; Lafkas et al., 2015; Marcet et al., 2011; Song et al., 2014; Tsao et al., 2009; Zhou et al., 2015). This is followed by the activation of a transcriptional program by the Geminin family members GEMC1 (*GMNC*) and MCIDAS/Multicilin (*MCIDAS*) that interact with E2F4/5-DP1 and are required for the generation of MCCs in frogs, fish and mammals (Arbi et al., 2016; Balestrini et al., 2010; Boon et al., 2014; Danielian et al., 2007; Kyrousi et al., 2015; Ma et al., 2014; Stubbs et al., 2012; Terre et al., 2016; Zhou et al., 2015). In addition, the transcription factors MYB(Pan et al., 2014; Tan et al., 2013), FOXJ1(You et al., 2004; Yu et al., 2008), RFX2/3(Chung et al., 2012; Didon et al., 2013; El Zein et al., 2009) and TAp73 (the transcriptionally active isoform encoded by the *Trp73* gene)(Marshall et al., 2016; Nemajerova et al., 2016), as well as the atypical cyclin, CCNO(Funk et al., 2015; Núnez-Ollé et al., 2017; Wallmeier et al., 2014), are necessary for the expression of genes that promote deuterosome mediated centriole amplification and the generation of multiple motile cilia. Mutations in human *MCIDAS* or *CCNO* underlie Reduced Generation of Multiple Motile Cilia (RGMC), a rare ciliopathy characterized by hydrocephaly, mucus accumulation and reduced fertility, all presumably due to defects in MCC differentiation(Amirav et al., 2016; Boon et al., 2014; Funk et al., 2015; Wallmeier et al., 2014).

Male and female infertility has been reported for a number of mice lacking genes involved in MCC development, including *Ccno*, the 3 *miR-34/449* genes, *FoxJ1*, *TAp73*, *E2F4 and 5* and *Gemc1*. Where this has been addressed in females, it appears to be due to the loss of MCCs in the oviducts (Chen et al., 1998; Marshall et al., 2016; Núnez-Ollé et al., 2017; Terre et al., 2016; Wu et al., 2014). However, in males, the origin of the defect has not been clearly established in most cases. Both the *miR-34/449* genes and *TAp73* are expressed in the testes and mice lacking the *miR-34/449* genes are impaired in meiosis and spermiogenesis and exhibit a “nearly empty” seminiferous tubule phenotype(Comazzetto et al., 2014; Wu et al., 2014; Yuan et al., 2015). *TAp73* deficient mice showed a similar seminiferous tubule phenotype(Tomasini et al., 2008), as well as degeneration of the Sertoli cells (SCs) that support spermatid development and ensure integrity of the blood-testes barrier(Holembowski et al., 2014). However, a conditional knockout of *E2f4* and *E2f5* alleles that targeted the EDs, but not spermatogonia or spermatocytes, also phenocopied the seminiferous tubule phenotype(Danielian et al., 2016), suggesting that defects in the EDs may be sufficient to strongly impair testes development.

Here we show that male mice lacking either *Gemc1* or *Ccno* exhibited a phenotype similar to mice lacking 3 members of the *Mir34/449* family, *TAp73* deletion or loss of both *E2f4* and *E2f5* (Comazzetto et al., 2014; Danielian et al., 2016; Holembowski et al., 2014; Inoue et al., 2014; Terre et al., 2016; Wu et al., 2014; Yuan et al., 2015). We found that mice lacking *Gemc1* or *Ccno* exhibited SC degeneration and a nearly empty seminiferous tubules phenotype. However, in mice lacking either gene, spermatids failed to enter the epididymis due to defects in MCC maturation. Moreover, we show that similar to *FoxJ1*, *Trp73* expression is high in the EDs and is dependent on GEMC1 but not CCNO, further establishing distinct temporal roles in the MCC transcriptional program. Our results demonstrate that GEMC1 and CCNO are required for ED MCC differentiation and suggest that these defects are likely to be the primary cause of male infertility in several mouse lines with MCC defects, and potentially in human RGMC patients.

## Results and Discussion

### GEMC1 is required for late stages of spermatogenesis

We analyzed adult testes of *Gemc1^-/-^* mice over the first three months and found no changes in size and weight compared with wild type (*Wt*) or *Gemc1^+/^-* littermates when normalized to body size (Fig. 1A). We next performed histological evaluation of testes during the first semi-synchronous wave of spermatogenesis (Fig. 1B). No overt differences were apparent between *Wt, Gemc1^+^/-* or *Gemc1^-/-^* mice during the first 20 days (Fig. 1C). However, by p27-p35, the thinning of the seminiferous epithelia became obvious, corresponding to the first appearance of elongating spermatids (ES) (Fig. 1D). Despite the onset of the seminiferous tubule phenotype, we observed similar numbers of mitotic cells, dead cells, normal meiotic progression and similar levels of hormonal expression in *Gemc1^-/-^* mice (Fig. S1A-E).

**Figure 1.**
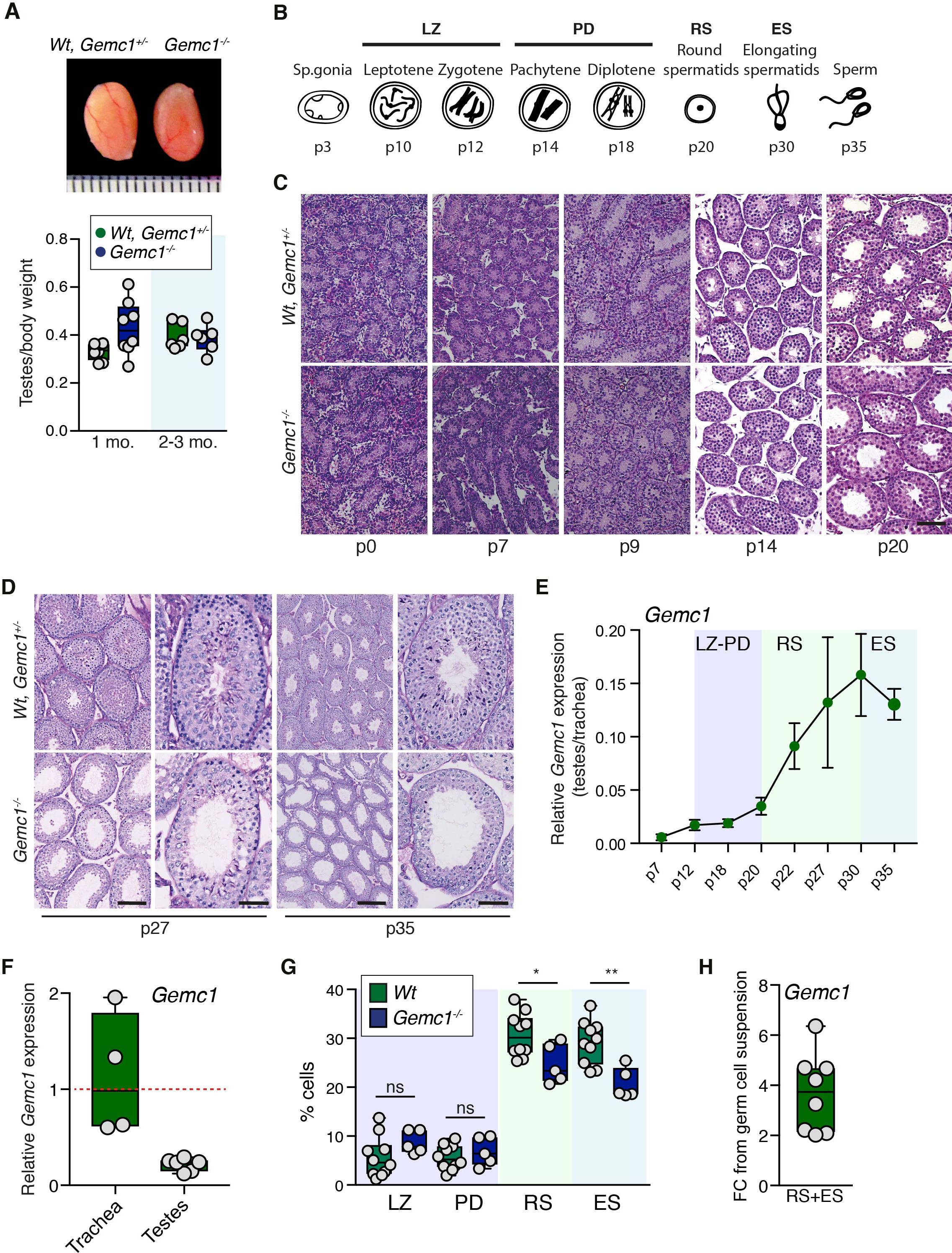
*Gemc1* loss impairs the late stages of spermatogenesis. **(A)** Example of dissected testes from littermate mice of the indicated genotypes at 3 months (top panel). Ruler=mm. Testes weight relative to the whole body at the indicated ages (n=8 for 1 month and n=6 animals per genotype for 2-3 month) is plotted below. (**B**) Schematic of semi-synchronous stages of spermatogenesis in mice (adapted from (Comazzetto et al., 2014)). (**C**) PAS staining of developing testes from *Wt*, *Gemc1^+/-^* and *Gemc1^-/-^* littermates at p0, p7, p9, p14 and p20. Scale bar=100μm. (**D**) PAS staining of p27 and p35 testes shows thinner seminiferous tubules and cell loss in the absence of *Gemc1*. Scale bars=100μm (left panels) and 50μm (right panels) for each age. (**E**) RT-PCR analysis of *Gemc1* expression from testicular RNA at the indicated postnatal days (n=2 for postnatal (p) days 7, 18, 20, 30 and n=3 animals for days 12, 22, 27, 35). Levels are plotted relative to that of the trachea and *Actb* was used as a normalization control. (**F**) RT-PCR analysis of *Gemc1* expression in 1-2 mo. old testes (n=6) compared to trachea (n=4). *Actb* was used as a normalization control. (**G**) Comparative abundance of each spermatogenic cell type of control and *Gemc1^-/-^* mice by fluorescence activated cell sorting (FACS) (*Wt* n=10, *Gemc1^-/-^* n=5 animals). *p=0.023 and **p=0.0023, unpaired t-test, two-tailed. (**H**) Ratio of *Gemc1* expression in isolated RS/ES populations compared to germ cell pellets determined by RTPCR (n=8 animals). *Actb* was used as a normalization control.

Consistent with our histological observations, *Gemc1* mRNA expression peaked around post-partum day 27 (p27), although overall levels were considerably lower than that in the trachea that contains a large number of MCCs (Figs. 1E,F). As the peak of *Gemc1* expression and appearance of tubule phenotypes correlated with late stages of spermatogenesis (Fig. 1B), we isolated and quantified enriched populations of testicular cell types (leptotene-zygotene (LZ), pachytene-diplotene (PD), round spermatids (RS) and elongating spermatids (ES)) by fluorescence activated cell sorting (FACS)(Fig. S1F). In testes from *Gemc1^-/-^* mice, we observed similar numbers of prophase cells (LZ and PD) but a significant reduction in RS and ES populations (Fig. 1G). This was further supported by the clear enrichment of *Gemc1* mRNA isolated from RS and ES populations compared to the germ cell pellet (Fig. 1H) and suggested that GEMC1 may support late stages of spermatogenesis, potentially through the control of transcriptional pathways.

### GEMC1 and CCNO prevent Sertoli cell degeneration

To address this possibility, we examined the transcriptional activation of key GEMC1 target genes in the testes. In contrast to tissues containing MCCs, we did not observe any significant alterations in several genes critical for MCC development, including *Ccno, Mcidas*, *FoxJ1 and Cdc20b* (Fig. 2A). Previous work reported a similar nearly empty seminiferous tubules phenotype in mice lacking *TAp73*, a gene recently linked to MCC formation, that was accompanied by extensive SC degeneration and spermatid detachment(Holembowski et al., 2014; Inoue et al., 2014; Marshall et al., 2016; Nemajerova et al., 2016; Tomasini et al., 2008). Examination of the seminiferous tubules of *Gemc1^-/-^* mice stained with Acetylated tubulin (Ac-tub) to visualize flagella revealed extensive spermatid detachment (Fig. 2B), suggesting similar defects in spermatid interactions with support cells may be present.

**Figure 2.**
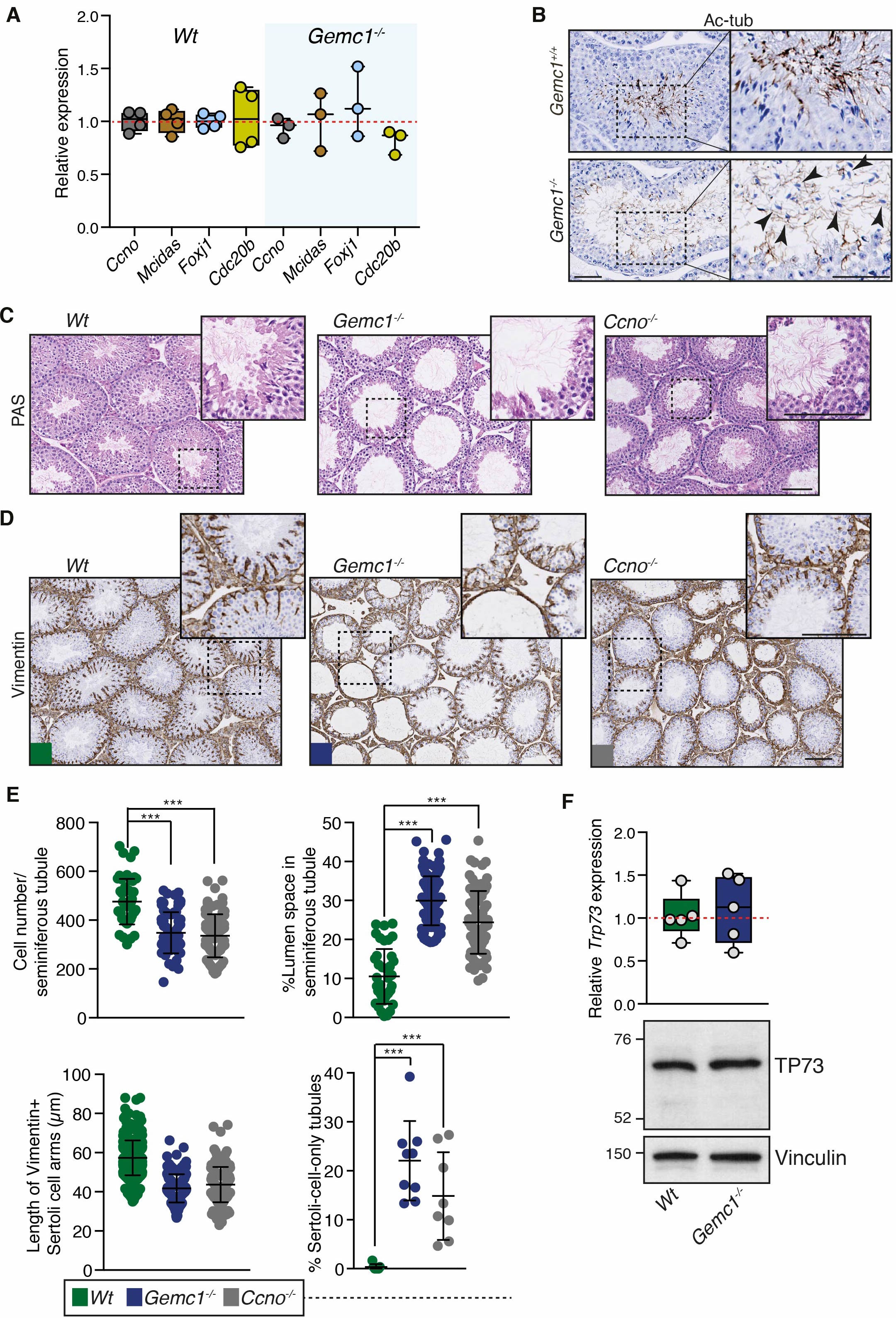
Gene expression and Sertoli cell degeneration in testes of mice lacking Gemc1 or Ccno. (**A**) RT-PCR analysis of the MCC targets *Ccno*, *Mcidas*, *FoxJ1* and *Cdc20b* in the p27 testes of *Wt* (n=4) and *Gemc1^-/-^* mice (n=3). (**B**) Ac-tubulin staining of the p35 seminiferous tubules of *Wt* and *Gemc1^-/-^* mice. Detached spermatids in *Gemc1^-/-^* indicated by black arrowheads. Scale bars=50μm. (**C**) PAS staining of testes sections from p35 *Gemc1^-/-^* and *Ccno^-/-^* mice revealed thinning of the spermatogenic cell layer. Scale bar=100μm. (**D**) Vimentin staining of SC intermediate filaments in the testes of the indicated genotype. Scale bars=250μm. (**E**) Quantifications of percent of cell number per seminiferous tubule (top left, n=4/genotype, ***p<0.0001), empty lumen space (n=4/genotype, top right, ***p<0.0001), length of vimentin positive SC arms (n=4, 2, 2 animals per genotype, bottom left) and SC only tubules (n=4/genotype, bottom right, ***p<0.0001 and p=0.0009). Results from p30-35 testes are shown; 10 tubules from two sections per animal were scored. Statistical analysis = unpaired t-test, two-tailed. (**F**) RT-PCR of *Trp73* mRNA levels (top panel, n=5 animals per genotype) and a representative western blot of TP73 levels in testes lysates from *Wt* or *Gemc1^-/-^* (n=2, bottom panel). *Actb* was used as a normalization control for mRNA levels and Vinculin was used as a loading control for protein levels.

As *Ccno^-/-^* males are also infertile(Núnez-Ollé et al., 2017), we histologically examined the testes of these animals in parallel to *Gemc1^-/-^*. We found that like *Gemc1^-/-^* mice, they exhibited a nearly empty seminiferous tubule phenotype (Fig. 1C). Immunostaining of Vimentin-containing intermediate filaments, one of the main components of the SC cytoskeleton (Aumuller et al., 1988), revealed marked structural abnormalities in the SCs of both *Gemc*1 and *Ccno* deficient testes (Fig. 1D). This was characterized by abnormally shorter and thinner cytoplasmic projections, as well as the appearance of SC-only seminiferous tubules (Fig.2C, D, E). In contrast, the SCs of *Wt* mice extended long cytoplasmic arms in contact with germ cell populations (Figs. 2C, D, E). Given the early transcriptional role of GEMC1 in MCCs, its expression during spermiogenesis and the strong similarity to the phenotypes reported for *TAp73* knockouts, we examined *Trp73* expression in the testes of *Gemc1^-/-^* mice. At both the mRNA and protein level, we saw no clear alteration in expression levels (Fig. 2F). These results showed that both *Gemc1* and *Ccno* phenocopied *TAp73* loss, exhibiting nearly empty seminiferous tubules and SC degeneration, but GEMC1 did not appear to influence the expression of its known targets or *Trp73* in testes.

### Defective efferent duct function in *Gemc1^-/-^* and *Ccno^-/-^* mice

Sperm enter the caput epididymis but remain immature until they reach the cauda epididymis, where they acquire motility and fertilization competency (Fig. 3A). *Gemc1^-/-^* and *Ccno^-/-^* epididymes appeared paler, smaller and thinner than *Wt* (Fig. 3B) and histological analysis revealed no detectable sperm in the caput, corpus or cauda epidiymes in either case, in contrast to *Wt* where sperm was abundant in all sections (Fig. 3B, C). Consistent with this, the lumen of both *Gemc1^-/-^* and *Ccno^-/-^* epididymes was often filled with PAS-positive material, a phenotype typically observed in the absence of spermatozoa(Abe and Takano, 1988). These results indicated that *Gemc1^-/-^* and *Ccno^-/-^* males have an apparently complete block in sperm transit to the epididymis.

**Figure 3.**
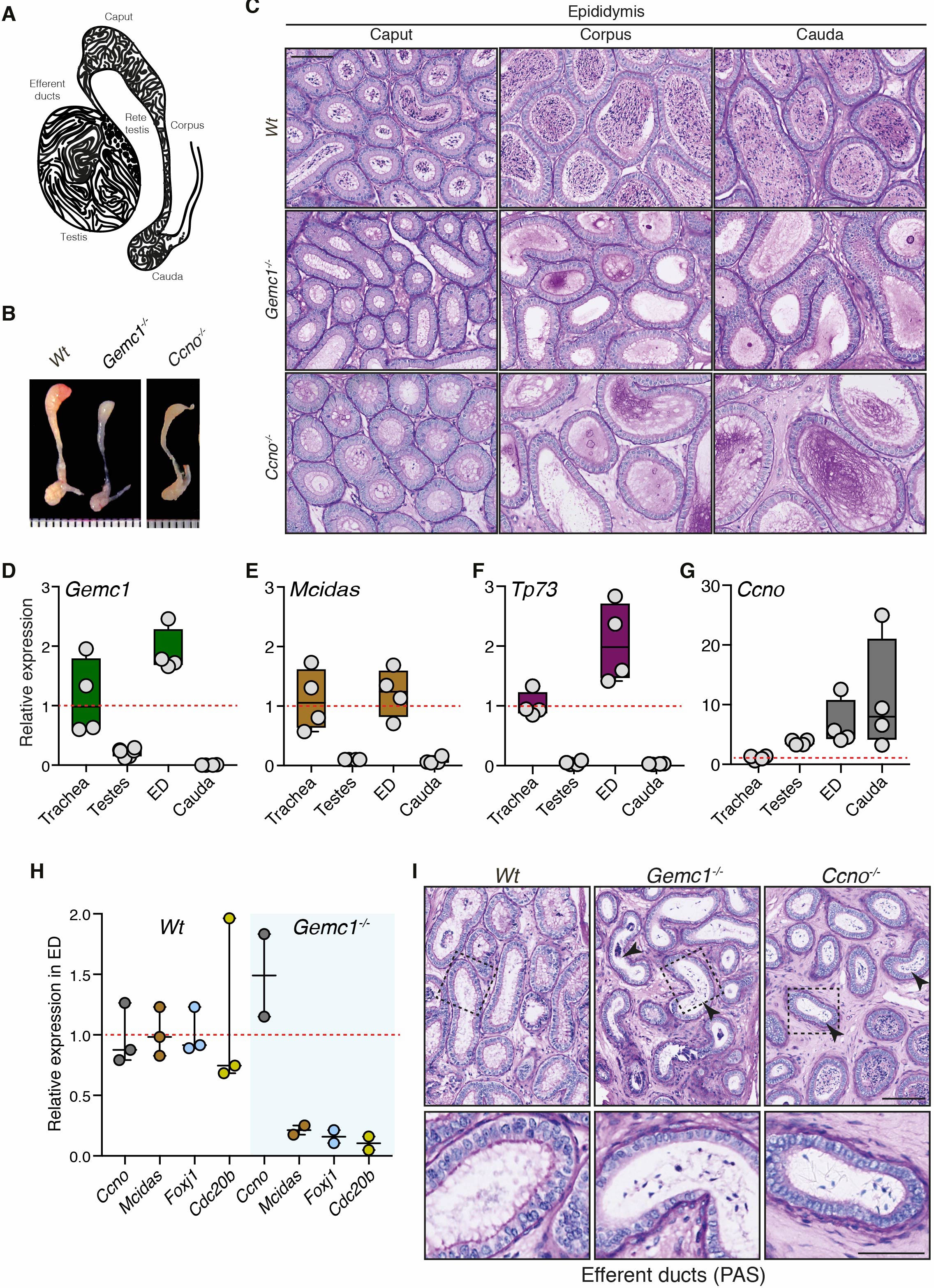
GEMC1 and CCNO are required for efferent duct formation. (**A**) Schematic of a vertical section of the testis, rete testes, efferent ducts and epididymis (caput, corpus, cauda). (**B**) Gross morphology of the epididymes of the indicated genotypes. Ruler=mm. (**C**). PAS staining of the three major regions of the mouse epididymis (caput, corpus and cauda) from adult mice of the indicated genotype. Scale bar=100μm and applies to all panels. (**D-G**) RTPCR analysis of the expression levels of the indicated gene in different tissues normalized to the trachea (n=4 animals). *Actb* was used as a normalization control. (**H**) RT-PCR analysis of the expression levels of the indicated gene in the efferent ducts of *Wt* (n=3) or *Gemc1^-/-^* animals (n=2). *Actb* was used as a normalization control. (**I**) PAS staining of the efferent ducts of mice of the indicated genotype. Black arrowheads indicate the accumulation of aberrant sperm in the *Gemc1^-/-^* and *Ccno^-/-^* mice compared to *Wt.* Scale bar=100μm (top panels) and 50μm (bottom panels).

As the EDs are critical for sperm transit into the epididymis and contain MCCs, we examined *Gemc1* and *Ccno* expression in the testes, cauda epididymis and EDs. The expression of *Gemc1* was considerably higher in the EDs than testes (Fig. 3D) and similar to its levels in the trachea that contains abundant multiciliated cells. A similar pattern was observed for its target gene *Mcidas* (Fig. 3E), as well as *Trp73* (Fig. 3F). In contrast, *Ccno* was only marginally upregulated in the ED compared to the testes and further increased in the cauda epididymis (Fig. 3G).

We next examined the expression levels of key MCC regulators in *Gemc1^-/-^* EDs. While *Ccno* levels were moderately increased, other known target genes including *Mcidas, Cdc20B*, and *FoxJ1*, as well as *Trp73*, were strongly downregulated (Fig. 3H), similar to what has been observed in the trachea(Terre et al., 2016). PAS staining of histological sections revealed that sperm accumulated in the EDs of both *Gemc1^-/-^* and *Ccno^-/-^* mice, where they are normally not detected in *Wt* animals due to the rapid transit through this region (Fig. 3I). Therefore, regardless of gene expression in the testes, spermatids were unable to enter the epidiymis of *Gemc1^-/-^* and *Ccno^-/-^* males due to defects in ED formation or function. In the case of GEMC1, this appeared to reflect its transcriptional role, similar to other MCC containing tissues.

### Distinct temporal roles of *Gemc1^-/-^* and *Ccno^-/-^* in the efferent ducts

Staining of motile cilia with acetylated tubulin (Ac-tub) revealed that MCCs were absent in *Gemc1^-/-^* EDs (Fig. 4A, top row middle panel) and a similar, although less severe, phenotype was observed in *Ccno^-/-^* mice (Fig. 4A, top row, right panel). In both mutants, Ac-tub staining was observed in trapped elongated spermatids in the central cavity of the ED.

**Figure 4.**
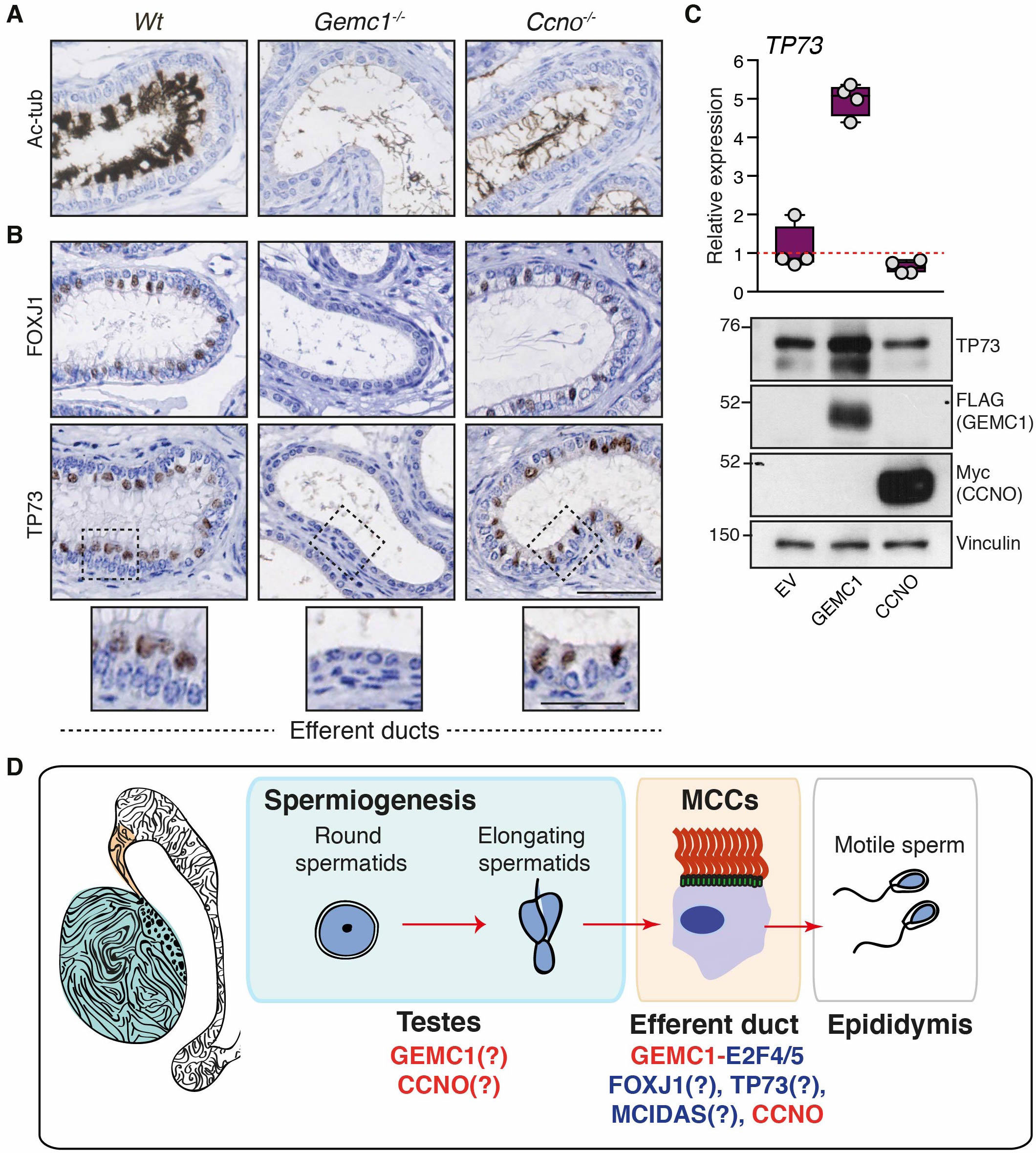
GEMC1 and CCNO are required for efferent duct development. (**A**) Representative Ac-tubulin staining of EDs of the indicated genotypes. Scale bars=100μm. (**B**) ED sections of the indicated genotypes stained with antibodies against FOXJ1 or TP73. Scale bars=100μm (main panels) and 50μm (bottom insets). (**C**) Overexpression of FLAG-GEMC1, but not Myc-CCNO, in AD293 cells by transient transfection induces TP73 expression. RTPCR was used to measure relative mRNA levels (n=4) and a representative western blot of 3 independent experiments is shown (bottom panel). *ACTB* was used as a normalization control. (**D**) Both *Gemc1* and *Ccno* are expressed in the testes and knockout mice exhibit strong phenotypes, including reduced cellularity and SC degeneration. However, both are expressed more highly in the ED and their loss leads to defective sperm transit, similar to what has been observed in E2F4/5 double mutants (Danielian et al., 2016). We propose that the failure of the EDs to generate fluid flow to support sperm transit is likely the primary cause of male infertility in several animals with MCC defects, and potentially human RGMC patients.

The ED epithelium of *Gemc1^-/-^* was thinner than both *Wt* and *Ccno^-/-^* mice (Fig. 4A), so we examined the expression of the FOXJ1 and TP73 transcription factors that are critical for MCC formation in other tissues(Marshall et al., 2016; Nemajerova et al., 2016; You et al., 2004). Cells lining the EDs were strongly immuno-positive for both proteins in *Wt* and *Ccno^-/-^* but staining was completely absent in *Gemc1^-/-^* (Fig. 4B, middle rows, bottom rows and insets). We have previously shown that the transient transfection of GEMC1 can activate early MCC factors, including both *MCIDAS* and *FOXJ1*, in AD293 cells(Terre et al., 2016). Transient overexpression of *GEMC1* led to a strong increase in *TP73* mRNA and protein levels, in contrast to *CCNO* expression that did not influence expression (Fig. 4C), demonstrating that GEMC1 is necessary and sufficient to activate TP73 expression. In contrast, CCNO plays a distinct role in ED MCC formation downstream of FOXJ1 and TP73 expression.

Collectively, our experiments have demonstrated that male mice lacking *Gemc1* or *Ccno* are infertile and exhibit a testes phenotype closely resembling that of *TAp73* deficient or *Mir449/34* triple knockout animals; a reduction in spermatocyte numbers, associated morphological aberrations, and degeneration of the SC support structures in testes (Comazzetto et al., 2014; Holembowski et al., 2014). However, despite generating some elongating spermatids, there is a complete failure of spermatids to enter the epididymis due to defects in MCCs. As the ED specific deletion of both *E2f4* and *E2f5*, the primary transcription factors involved in the MCC transcriptional cascade driven by GEMC1, also caused a similar nearly empty seminiferous tubule phenotype, this indicates that impairment of the MCC transcriptional program in the ED is likely sufficient to cause the phenotypes we have observed and we propose that it is most likely the dominant cause of infertility in *Gemc1^-/-^* and *Ccno^-/-^* mice, and potentially in male RGMC patients (Fig. 4D).

While the role of GEMC1 appears to be clearly related to transcriptional activation, the function of CCNO remains enigmatic. Recent work demonstrated that the mitotic oscillator machinery and the fine tuning of CDK1 activity is required for the stepwise development and dissociation of deuterosomes in mouse neuronal progenitors (Al Jord et al., 2017). Given that CCNO interacts with CDK1(Roig et al., 2009), it seems plausible that one of its primary functions may be the regulation of CDK1 activity during deuterosome formation. Future work will be needed to understand precisely how GEMC1 and CCNO regulate different aspects of the transcriptional response and deuterosome formation, as well as their potential roles in other tissues.

## Materials and Methods

### Histopathology and immunohistochemistry of murine tissues

*Gemc1^-/-^* and *Ccno^-/-^* mice on a mixed C57BL/6-129SvEv background were described previously (Núnez-Ollé et al., 2017; Terre et al., 2016). Animals were maintained in accordance with the European Community (86/609/EEC) guidelines in the Specific-Pathogen Free (SPF) facilities of the Barcelona Science Park (PCB). Protocols were approved by the Animal Care and Use Committee of the PCB (IACUC; CEEA-PCB) in accordance with applicable legislation (Law 5/1995/GC; Order 214/1997/GC; Law 1201/2005/SG). All efforts were made to minimize use and suffering. Sample sizes were not defined to detect pre-determined effect size. Animals were not randomized, were identified by genotyping for analysis and ages, were all males and ages are indicated in the figure legends. Testes and epididymis were harvested and fixed in 4% PFA or Bouin’s solution (Electron Microscopy Sciences) overnight at 4 ºC and embedded in paraffin using standard procedures. Sections were cut at 10 μm thickness and stained with hematoxylin and eosin (H&E) and Periodic acid/Schiff reagent (PAS, Sigma-Aldrich). For colorimetric visualization, sections were incubated with primary antibody overnight at RT after quenching endogenous peroxidase using 0.6% H_2_O_2_ (vol/vol) in methanol. Slides were washed and incubated with biotinylated secondary antibody and avidin-biotin complex (Vectastain Elite kit, Vector Labs). Immunoreactive signals were visualized with the VIP substrate kit (Vector Labs) using the manufacturer’s protocol. Sections were counterstained with 0.1% methyl green (wt/vol), dehydrated, and mounted in DPX (Fluka).

### Antibodies

Primary antibodies used in the publication:

| Antigen | Species | Vendor | Clone | Reference no. | Dilution |
| --- | --- | --- | --- | --- | --- |
| Ac-tub | mouse | Sigma | 6-11B-1 | T6793 | 1:1000 |
| Vimentin | rabbit | Abcam | EPR3776 | ab92547 | 1:2000 |
| Vinculin | rabbit | Abcam | EPR8185 | ab129002 | 1:2000 |
| FOXJ1 | mouse | eBioscience | 2A5 | 14-9965-83 | 1:500 |
| TP73 | rabbit | Abcam | EP436Y | ab40658 | 1:50 |
| Myc | rabbit | Abcam |  | ab9132 | 1:1000 |
| FLAG | mouse | Sigma | M2 | F3165 | 1:1000 |

All antibodies have been previously validated and were subjected to additional controls. The Ac-tub antibody has been used widely in the field and recognizes multiciliated cells that are absent in *Gemc1* deficient mice(Terre et al., 2016). The FOXJ1 and Vimentin antibodies have been validated in knockouts and extensively in publications (see Thermofisher and Abcam product page) and the TP73 was validated in knockout animals(Marshall et al., 2016) and signal is in the expected cell types and correlates with the mRNA level in our experiments. The Vinculin antibody recognizes a band of the correct size and was used as a loading control. Myc and FLAG epitope antibodies have been validated extensively and recognize proteins of the correct size only when tagged cDNAs are expressed.

### Microscopy

Histological sections from testes, epididymis and efferent ducts were imaged with the digital slide scanner Nanozoomer 2.0HT (Hamamatsu) and analyzed using the NDP view 2 free software (Hamamatsu). Image analysis and quantification of IHC were performed with the TMARKER free software (GitHub). For quantitation of cell numbers per tubule, ten random tubules from two PAS-stained sections were counted. For quantitation of % of lumen space, tubule area and lumen area were analyzed for ten tubules of PAS-stained sections (two sections per animal). The length of cytoplasmic Sertoli cell arms was determined in μm using Vimentin staining. Vimentin marks intermediate filaments that are seen as cytoplasmic arms surrounding the nucleus and extending from the basal region towards the tubular lumen. Sertoli cell arm’s length was calculated in ten tubules of Vimentin-stained sections (2 sections per animal). Vimentin was also used to detect Sertoli-cell-only tubules, characterized by intense staining of the entire seminiferous tubule combined with the lack or dramatic decrease of germ cells.

### Quantitative real-time PCR (qRT-PCR)

Dissected testes, epididymis, efferent ducts or pituitary gland were carefully dissected and collected on ice, washed in PBS and frozen. Testes were disrupted in Tri-Reagent (Sigma) by zirconium beads in a mechanical tissue disruptor (Precellys 24, Bertin technologies). Total RNA was isolated according to manufacturer recommendations (PureLink RNA mini kit, Ambion) and 1ug of RNA was treated with DNase I prior to cDNA synthesis (Thermo Fisher). cDNA was generated using 0.5-1 μg of total RNA and a High Capacity RNA-to-cDNA Kit (Applied Biosystems). Quantitative real time-PCR (RT-QPCR) was performed using the comparative CT method and a Step-One-Plus real-time PCR Applied Biosystems Instrument. Amplification was performed using Power SYBR Green PCR Master Mix (Applied Biosystems) or TaqMan Universal PCR Master Mix (Applied Biosystems). All assays were performed in duplicate. For TaqMan assays, *ActB* (mouse and human) probe was used as an endogenous control for normalization and a specific Taqman probe was used for mouse or human *Gemc1*(*Mm02581229_m1), Mcidas (Mm01308202_m1), Ccno(Mm01297259_m1), FoxJ1 (Mm01267279_m1)* and *Trp73 (Mm00660220_m1) and TP73 (Hs01056231_m1).* Primers used for SYBR Green assays (Sigma) are listed below.

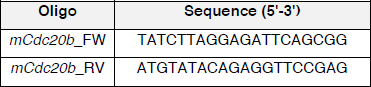

### Germ cell isolation for fluorescence-activated cell sorting (FACS)

Testes from adult male mice were isolated, decapsulated and processed together in a 15 ml falcon tube. Testes were first digested in EKRB medium (120 mM NaCl, 4.8 mM KCl, 25.2 mM NaHCO_3_, 1.2 mM KH_2_PO_2_, 1.2 mM MgSO4, 1.3 mM CaCl2, 11 mM Glucose, non-essential aminoacid (Invitrogen), penicillin-streptomycin (Invitrogen)) with collagenase (0.5 mg/ml, Sigma T6763) in shaking water bath at 32 ºC for 10 min. Seminiferous tubules were let sediment by gravity for 1-3 min and washed twice with EKRB. The tubules were further dissociated in EKRB with trypsin (1 mg/ml, Sigma T6763) and DNase (5 μg/ml, Sigma D4263) in shaking at 32 ºC for 15 min. Cells were resuspended thoroughly with a pipette until obtaining a single cell suspension and 1 ml of FBS was added to neutralize the Trypsin. Cell suspension was filtered with a 70 μm cell strainer and total cell number was counted using the Neubauer chamber. 1 million cells/ml were resuspended in EKRB supplemented with 10% FBS and Hoechst 33342 (10 μg/ml) (Life Technologies) in shaking at 32 ºC for 30-60 min. Propidium iodide (PI, 30 μg/ml) (Life Technologies) was added for the discrimination of dead cells. Flow cytometric experiments were carried out using a FacsAria I SORP cell sorter (Beckton Dickinson, San Jose, California), using a 70-micron nozzle at 60PSI. Excitation of the sample was done using a blue (488nm) laser for forward scatter (FSC) parameter; green-orange laser (561nm) was used for the excitation of PI and side scatter (SSC) signal, and a UV laser (350nm) was used for Hoechst 33342 excitation. Cells were gated according to their scatter (FSC vs SSC) parameters; fluorescence of Hoechst 33342 was measured on live (not stained with PI), non-aggregated cells. Red emission (660/40 nm) vs. blue emission (395/25 nm) from the UV laser was used on a dot plot in order to discriminate populations. Results were analyzed using the FlowJo software.

### Immunofluorescence

Slides containing meiotic squashes were then washed several times in PBS followed with washing in PBS-T (0.4% Triton X-100 in PBS) and blocked with 5% goat serum and 1% BSA in PBS-T for 1 h. Slides were incubated overnight at 4 ºC with primary antibody, washed several times in PBS-T and stained with the Alexa Fluor-conjugated complementary antibody (Life Technologies) for 1 h at RT. After final washing, DNA was counterstained with DAPI. Slides were mounted using Vectashield Antifade reagent (Vector Laboratories) and imaged using a Leica TCS SP5 confocal microscope equipped with 63x NA 1.40 oil immersion objective and HyD detectors.

### Cell culture, transfection and Western Blotting

AD293 cells (Stratagene) were cultured in DMEM (Gibco) with 10% FBS (Hyclone) and routinely tested for mycoplasma and found negative. For transient transfections, AD293 cells were seeded in 10 cm plates at 70% confluence and 10μg of plasmid were transfected the day after with Polyethylenimine (Polysciences). The medium was changed 12 h post-transfection and cells were collected 48 h after with RIPA buffer (50mM TrisHCl pH 8, 150mM NaCl, 1% NP-40, 0.1% SDS, 0.5% Sodium Deoxycholate). For tissue samples, testes were disrupted in RIPA buffer using zirconium beads in a mechanical tissue disruptor (Precellys 24, Bertin technologies). Samples were incubated 20 min on ice and sonicated using a Bioruptor XL sonication device (Diagenode) for 15 min with 15 s intervals and centrifuged at 4 ºC for 20 min at 1300 rpm. Antibodies to the following epitope tags were used: Flag-tag (Sigma, F3165), Myc-tag (Abcam, ab9132), p73 EP436Y (Abcam, ab50658), and Vinculin (Sigma, V9264).

### DNA constructs

The expression construct for FLAG tagged human *GEMC1* (pcDNA5-FRT/TO-flag-hGemc1) was previously described in(Terre et al., 2016), Human *CCNO* cDNA was obtained from Expressed Sequence Tag EST IMAGE: 6421733 (gi22331975) as a template and amplified using 5’- GCGAATTCCATGGTGACCCCCTGTCCCACCAGCC-3’ and 5’- GCTCTAGATTATTTCGAGCT CGGGGGCAGG-3’ primers. hCCNO cDNA was cloned into the pBluescript (I) SK(+) vector (Addgene) at the EcoRI and XbaI restriction sites and then into the mammalian expression vector pCDNA3.1 (Invitrogen) modified with a n N-terminal myc-tag to produce proteins N-terminally fused to myc under the control of the constitutive CMV promoter (pcDNA3.1-myc-hCcno). The pCMX-flag plasmid was used as an empty-vector control (a gift from Ron Evans, the Salk Institute for Biological Studies).

## Acknowledgements

We are grateful to Haotian Zhao, Muriel Lize, Heymut Omran and Julia Wallmeier for discussing unpublished data and experimental suggestions, to Monica Di Giacomo and Ignasi Roig for helpful input, Lluis Palenzuela for mouse colony management, Neus Prats and the IRB Barcelona histology facility for assistance with IHC and protocol establishment and Oscar Reina of the IRB Biostatistics facility for help with data analysis.

## Funding

THS is funded by the Spanish Ministry of Economy and Competitiveness (MINECO) (BFU2015-68354, Ayudas para incentivar la incorporación estable de doctores (IED) 2015)) and institutional funding from MINECO through the Centres of Excellence Severo Ochoa award and from the CERCA Programme of the Catalan Government. BT and ML are supported by Severo Ochoa FPI fellowships (MINECO) and GG is supported by ISCIII-grant PI13/00864 and FEDER Funds.

## Competing interests

The authors declare no competing interests

## References

Abe, K. and Takano, H. (1988). Changes in distribution and staining reactivity of PAS-positive material in the mouse epididymal duct after efferent duct ligation. Arch Histol Cytol 51, 433–441.

Al Jord, A., Shihavuddin, A., Servignat d’Aout, R., Faucourt, M., Genovesio, A., Karaiskou, A., Sobczak-Thepot, J., Spassky, N. and Meunier, A. (2017). Calibrated mitotic oscillator drives motile ciliogenesis. Science 358, 803–806.

Amirav, I., Wallmeier, J., Loges, N. T., Menchen, T., Pennekamp, P., Mussaffi, H. Abitbul, R., Avital, A., Bentur, L., Dougherty, G. W., et al. (2016). Systematic Analysis of CCNO Variants in a Defined Population: Implications for Clinical Phenotype and Differential Diagnosis. Hum Mutat 37, 396–405.

Arbi, M., Pefani, D. E., Kyrousi, C., Lalioti, M. E., Kalogeropoulou, A., Papanastasiou, A. D., Taraviras, S. and Lygerou, Z. (2016). GemC1 controls multiciliogenesis in the airway epithelium. EMBO Rep 17, 400–413.

Aumuller, G., Steinbruck, M., Krause, W. and Wagner, H. J. (1988). Distribution of vimentin-type intermediate filaments in Sertoli cells of the human testis, normal and pathologic. Anat Embryol (Berl) 178, 129–136.

Balestrini, A., Cosentino, C., Errico, A., Garner, E. and Costanzo, V. (2010). GEMC1 is a TopBP1-interacting protein required for chromosomal DNA replication. Nat Cell Biol 12, 484–491.

Boon, M., Wallmeier, J., Ma, L., Loges, N. T., Jaspers, M., Olbrich, H., Dougherty, G. W., Raidt, J., Werner, C., Amirav, I., et al. (2014). MCIDAS mutations result in a mucociliary clearance disorder with reduced generation of multiple motile cilia. Nature communications 5, 4418.

Chen, J., Knowles, H. J., Hebert, J. L. and Hackett, B. P. (1998). Mutation of the mouse hepatocyte nuclear factor/forkhead homologue 4 gene results in an absence of cilia and random left-right asymmetry. J Clin Invest 102, 1077–1082.

Chung, M. I., Peyrot, S. M., LeBoeuf, S., Park, T. J., McGary, K. L., Marcotte, E. M. and Wallingford, J. B. (2012). RFX2 is broadly required for ciliogenesis during vertebrate development. Dev Biol 363, 155–165.

Comazzetto, S., Di Giacomo, M., Rasmussen, K. D., Much, C., Azzi, C., Perlas, E., Morgan, M. and O’Carroll, D. (2014). Oligoasthenoteratozoospermia and infertility in mice deficient for miR-34b/c and miR-449 loci. PLoS Genet 10, e1004597.

Danielian, P. S., Bender Kim, C. F., Caron, A. M., Vasile, E., Bronson, R. T. and Lees, J. A. (2007). E2f4 is required for normal development of the airway epithelium. Dev Biol 305, 564–576.

Danielian, P. S., Hess, R. A. and Lees, J. A. (2016). E2f4 and E2f5 are essential for the development of the male reproductive system. Cell Cycle 15, 250–260.

Didon, L., Zwick, R. K., Chao, I. W., Walters, M. S., Wang, R., Hackett, N. R. and Crystal, R. G. (2013). RFX3 modulation of FOXJ1 regulation of cilia genes in the human airway epithelium. Respiratory research 14, 70.

El Zein, L., Ait-Lounis, A., Morle, L., Thomas, J., Chhin, B., Spassky, N., Reith, W. and Durand, B. (2009). RFX3 governs growth and beating efficiency of motile cilia in mouse and controls the expression of genes involved in human ciliopathies. J Cell Sci 122, 3180–3189.

Funk, M. C., Bera, A. N., Menchen, T., Kuales, G., Thriene, K., Lienkamp, S. S., Dengjel, J., Omran, H., Frank, M. and Arnold, S. J. (2015). Cyclin O (Ccno) functions during deuterosome-mediated centriole amplification of multiciliated cells. EMBO J

Holembowski, L., Kramer, D., Riedel, D., Sordella, R., Nemajerova, A., Dobbelstein, M. and Moll, U. M. (2014). TAp73 is essential for germ cell adhesion and maturation in testis. J Cell Biol 204, 1173–1190.

Inoue, S., Tomasini, R., Rufini, A., Elia, A. J., Agostini, M., Amelio, I., Cescon, D., Dinsdale, D., Zhou, L., Harris, I. S., et al. (2014). TAp73 is required for spermatogenesis and the maintenance of male fertility. Proc Natl Acad Sci U S A 111, 1843–1848.

Joseph, A., Shur, B. D. and Hess, R. A. (2011). Estrogen, efferent ductules, and the epididymis. Biol Reprod 84, 207–217.

Kyrousi, C., Arbi, M., Pilz, G. A., Pefani, D. E., Lalioti, M. E., Ninkovic, J., Gotz, M., Lygerou, Z. and Taraviras, S. (2015). Mcidas and GemC1/Lynkeas are key regulators for the generation of multiciliated ependymal cells in the adult neurogenic niche. Development.

Lafkas, D., Shelton, A., Chiu, C., de Leon Boenig, G., Chen, Y., Stawicki, S. S., Siltanen, C., Reichelt, M., Zhou, M., Wu, X., et al. (2015). Therapeutic antibodies reveal Notch control of transdifferentiation in the adult lung. Nature.

Ma, L., Quigley, I., Omran, H. and Kintner, C. (2014). Multicilin drives centriole biogenesis via E2f proteins. Genes Dev 28, 1461–1471.

Marcet, B., Chevalier, B., Luxardi, G., Coraux, C., Zaragosi, L. E., Cibois, M., Robbe-Sermesant, K., Jolly, T., Cardinaud, B., Moreilhon, C., et al. (2011). Control of vertebrate multiciliogenesis by miR-449 through direct repression of the Delta/Notch pathway. Nat Cell Biol 13, 693–699.

Marshall, C. B., Mays, D. J., Beeler, J. S., Rosenbluth, J. M., Boyd, K. L., Santos Guasch, G. L., Shaver, T. M., Tang, L. J., Liu, Q., Shyr, Y., et al. (2016). p73 Is Required for Multiciliogenesis and Regulates the Foxj1-Associated Gene Network. Cell Rep 14, 2289–2300.

Nemajerova, A., Kramer, D., Siller, S. S., Herr, C., Shomroni, O., Pena, T., Gallinas Suazo, C., Glaser, K., Wildung, M., Steffen, H., et al. (2016). TAp73 is a central transcriptional regulator of airway multiciliogenesis. Genes Dev 30, 1300–1312.

Núnez-Ollé, M., Jung, C., Terré, B., Balsiger, N. A., Plata, C., Roset, R., Pardo-Pastor, C., Garrido, M., Rojas, S., Alameda, F., et al. (2017). Constitutive Cyclin O deficiency results in penetrant hydrocephalus, impaired growth and infertility. Oncotarget 8, 99261–99273.

Pan, J. H., Adair-Kirk, T. L., Patel, A. C., Huang, T., Yozamp, N. S., Xu, J., Reddy, E. P., Byers, D. E., Pierce, R. A., Holtzman, M. J., et al. (2014). Myb permits multilineage airway epithelial cell differentiation. Stem Cells 32, 3245–3256.

Roig, M. B., Roset, R., Ortet, L., Balsiger, N. A., Anfosso, A., Cabellos, L., Garrido, M., Alameda, F., Brady, H. J. and Gil-Gomez, G. (2009). Identification of a novel cyclin required for the intrinsic apoptosis pathway in lymphoid cells. Cell Death Differ 16, 230–243.

Song, R., Walentek, P., Sponer, N., Klimke, A., Lee, J. S., Dixon, G., Harland, R., Wan, Y., Lishko, P., Lize, M., et al. (2014). miR-34/449 miRNAs are required for motile ciliogenesis by repressing cp110. Nature 510, 115–120.

Steger, K. (2001). Haploid spermatids exhibit translationally repressed mRNAs. Anat Embryol (Berl) 203, 323–334.

Stubbs, J. L., Vladar, E. K., Axelrod, J. D. and Kintner, C. (2012). Multicilin promotes centriole assembly and ciliogenesis during multiciliate cell differentiation. Nat Cell Biol 14, 140–147.

Tan, F. E., Vladar, E. K., Ma, L., Fuentealba, L. C., Hoh, R., Espinoza, F. H., Axelrod, J. D., Alvarez-Buylla, A., Stearns, T., Kintner, C., et al. (2013). Myb promotes centriole amplification and later steps of the multiciliogenesis program. Development 140, 4277–4286.

Terre, B., Piergiovanni, G., Segura-Bayona, S., Gil-Gomez, G., Youssef, S. A., Attolini, C. S., Wilsch-Brauninger, M., Jung, C., Rojas, A. M., Marjanovic, M., et al. (2016). GEMC1 is a critical regulator of multiciliated cell differentiation. EMBO J 35, 942–960.

Tomasini, R., Tsuchihara, K., Wilhelm, M., Fujitani, M., Rufini, A., Cheung, C. C., Khan, F., Itie-Youten, A., Wakeham, A., Tsao, M. S., et al. (2008). TAp73 knockout shows genomic instability with infertility and tumor suppressor functions. Genes Dev 22, 2677–2691.

Tsao, P. N., Vasconcelos, M., Izvolsky, K. I., Qian, J., Lu, J. and Cardoso, W. V. (2009). Notch signaling controls the balance of ciliated and secretory cell fates in developing airways. Development 136, 2297–2307.

Wallmeier, J., Al-Mutairi, D. A., Chen, C. T., Loges, N. T., Pennekamp, P., Menchen, T., Ma, L., Shamseldin, H. E., Olbrich, H., Dougherty, G. W., et al. (2014). Mutations in CCNO result in congenital mucociliary clearance disorder with reduced generation of multiple motile cilia. Nat Genet 46, 646–651.

Wu, J., Bao, J., Kim, M., Yuan, S., Tang, C., Zheng, H., Mastick, G. S., Xu, C. and Yan, W. (2014). Two miRNA clusters, miR-34b/c and miR-449, are essential for normal brain development, motile ciliogenesis, and spermatogenesis. Proc Natl Acad Sci U S A 111, E2851–2857.

You, Y., Huang, T., Richer, E. J., Schmidt, J. E., Zabner, J., Borok, Z. and Brody, S. L. (2004). Role of f-box factor foxj1 in differentiation of ciliated airway epithelial cells. American journal of physiology. Lung cellular and molecular physiology 286, L650–657.

Yu, X., Ng, C. P., Habacher, H. and Roy, S. (2008). Foxj1 transcription factors are master regulators of the motile ciliogenic program. Nat Genet 40, 1445–1453.

Yuan, S., Tang, C., Zhang, Y., Wu, J., Bao, J., Zheng, H., Xu, C. and Yan, W. (2015). mir-34b/c and mir-449a/b/c are required for spermatogenesis, but not for the first cleavage division in mice. Biology open 4, 212–223.

Zhou, F., Narasimhan, V., Shboul, M., Chong, Y. L., Reversade, B. and Roy, S. (2015). Gmnc Is a Master Regulator of the Multiciliated Cell Differentiation Program. Curr Biol 25, 3267–3273.

